# The effects of long-term exercise training on the neural control of walking

**DOI:** 10.1101/2021.01.21.427603

**Authors:** Morteza Yaserifar, Ziya Fallah Mohammadi, Sayed Esmaiel Hosseininejad, Iman Esmaili Paeen Afrakoti, Kenneth Meijer, Tjeerd W. Boonstra

**Affiliations:** Department of Neuropsychology & Psychopharmacology, Faculty of Psychology and Neuroscience, Maastricht University, Maastricht, Netherland; Department of Exercise Physiology, Faculty of Physical Education and Sport Science, University of Mazandaran, Mazandaran, Iran; Department of Sports Biomechanics, Faculty of Physical Education and Sport Science, University of Mazandaran, Mazandaran, Iran; Department of Electrical Engineering, Faculty of Technology and Engineering, University of Mazandaran, Mazandaran, Iran; Department of Nutrition and Movement Sciences, NUTRIM School for Nutrition and Translational Research in Metabolism, Faculty of Health, Medicine and Life Sciences, Maastricht University, Maastricht, Netherland; Neuroscience Research Australia, Sydney, Australia

**Author notes:** **Corresponding Author:** Dr Tjeerd W. Boonstra, Department Neuropsychology & Psychopharmacology, Faculty of Psychology and Neuroscience, Maastricht University, Maastricht, Netherland.

**Keywords:** Walking, Electromyography, Muscle synergy, Motor control, Exercise training, Gait parameters

## Abstract

How does long-term training modify the neural control of walking? Here we investigate changes in kinematics and muscle synergies of the lower extremities in 10 soccer players and 10 non-athletes while they walked with eyes open or closed either overground or on a treadmill. Electromyography (EMG) was acquired from eight muscles of the right leg and foot switch data were recorded to extract temporal gait parameters. Muscle synergies were extracted using non-negative matrix factorisation for each participant and condition separately and were then grouped using k-means clustering. We found that both the cycle and stance duration were longer during treadmill walking compared to overground walking, whereas the swing phase was longer during the eyes-open compare to the eyes-closed condition. On average, more synergies were expressed in the athlete compared to the non-athlete group and during treadmill compared to overground walking. We found that synergy 2 involved in ankle plantarflexion was more often activated in athletes than in non-athletes. We did not find statistical group differences for the synergy metrics but several differences were observed between conditions: peak activation of synergy 5 (VM and VL muscles) increased during overground walking compared to treadmill walking. In addition, reduced activation of synergy 3 (TA muscle) and synergy 4 was observed during eyes-closed compared to eyes-open walking. These findings suggest that during walking long-term training results in greater flexibility of muscle coordination by recruiting additional synergies, but we found no evidence that long-term training affects the activation patterns of these synergies.

## 1. Introduction

Walking is a daily activity that we perform with little effort or attention, but requires delicate control of our complex musculoskeletal system (Aoi et al., 2019; Nielsen, 2003). Despite of the many degrees of freedom of the musculoskeletal system and the complexity human gait, gait patterns tend to be stereotypical and have close to optimal energy expenditure (Ackermann & Van den Bogert, 2010; Clever, Schemschat, Felis, & Mombaur, 2016). This means that out of infinitive options, the central nervous system (CNS) selects a smooth and coordinated movement trajectory that minimizes joint torques or fatigue. However, what an optimal gait trajectory is depends on characteristics of the individual and the surrounding and considerable variability can hence be observed across gait cycles, individuals and task conditions (Hausdorff, 2005; Kang & Dingwell, 2008; Springer et al., 2006). For example, the neuromuscular control of walking is affected by physical health, revealing changes in gait and balance across the lifespan (Voelcker-Rehage & Niemann, 2013) and with exercise training (Godde & Voelcker-Rehage, 2017).

Research on exercise training has mostly focused on the reduction of injuries through reinforcing muscle groups, emphasizing a combination of strength, balance and aerobic training (Sherrington et al., 2008). For instance, strength and balance training can reduce fall risk during walking (McCrum, Gerards, Karamanidis, Zijlstra, & Meijer, 2017). Also, people in the early stage of Parkinson disease show increased gait speed, step and stride length, and hip and ankle joint excursion during gait training and also improved weight distribution in the sit-to-stand test after completing 24 exercise sessions over 8 weeks (Fisher et al., 2008). Furthermore, patients who had experienced stroke improved walking speed, stride length and symmetry index after three weeks of backward walking training (Yang, Yen, Wang, Yen, & Lieu, 2005). Finally, treadmill training with visual feedback showed the efficiency of training program in patients in the late period after stroke: Participants had a great improvement in shortening of the stance phase, lengthening of the swing phase, and increasing cycle length in the unaffected limb after two weeks treadmill training (Drużbicki, Guzik, Przysada, Kwolek, & Brzozowska-Magoń, 2015).

Although the behavioural improvements of exercise training are extensively described, the physiological changes underlying these improvements in performance are less well understood. Several changes in the central nervous system have been reported. For example, exercise training can enhance neurotrophic factors (Perrey, 2013) and increase interhemispheric brain activity at the cortical level (Lepley, Lepley, Onate, & Grooms, 2017). However, it remains unclear how these changes in the CNS relate to the improvements observed at the behavioural level. Muscle synergy analysis is a well-establish approach to investigate the neural coordination of movement (d’Avella & Bizzi, 2005; Torres-Oviedo & Ting, 2007; Tresch & Jarc, 2009) and has been used to investigate changes in control strategy resulting from exercise training (Safavynia, Torres-Oviedo, & Ting, 2011). Muscle synergies are defined as patterns of muscle co-activation that are activated by single neural command signal (Nordin & Dufek, 2016) and have been quantified using non-negative matrix factorisation of electromyography acquired from multiple muscles (Tresch, Cheung, & d’Avella, 2006). During unimpaired gait, less than six synergies are sufficient to describe the muscle activities during walking (Allen & Neptune, 2012; Chvatal & Ting, 2013; De Groote, Jonkers, & Duysens, 2014; Ivanenko, Cappellini, Dominici, Poppele, & Lacquaniti, 2005). Each synergy involves a specific group of muscles that is activated during a specific phase of the gait cycle (Ivanenko, Poppele, & Lacquaniti, 2006). For example, knee extensor groups provide body support in early stance while ankle plantar flexors are more activated in the late stance phase (Allen & Neptune, 2012; Ivanenko et al., 2005; A. S. Oliveira, Gizzi, Farina, & Kersting, 2014).

Some evidence indicates that muscle synergies can indeed detect changes in control strategy with improvement or decline of performance. For example, when muscle synergies were extracted during overground walking and the beam-walking test, experts used more synergies during beam-walking, possibly to create greater efficiency in muscle activity by transferring the learned muscle activity patterns to related tasks (Sawers, Allen, & Ting, 2015). Furthermore, it has been reported that patients use fewer muscle synergies than healthy subjects during walking (Clark, Ting, Zajac, & Neptune, 2009; A. S. C. Oliveira, Gizzi, Kersting, & Farina, 2012; Pérez-Nombela et al., 2017). The increase in the number of muscle synergies can reduce muscle coactivity when performing the same task (Sawers et al., 2015). Furthermore, balance investigation between athletes and non-athletes showed that despite using the same number of muscle synergy, athletes used a low co-activation strategy of ankle stabilizer muscles during balance task (M. Kim, Kim, Kim, & Yoon, 2018). Applying functional electrical stimulation training on patients who just could walk showed a general improvement in muscle coordination with an increase in the number of synergies (Ferrante et al., 2016). These findings suggest that the CNS may change muscle coordination either by increasing the number of muscle synergies or reducing co-activation.

Here, we investigate differences in muscle synergies during walking in athletes and non-athletes to examine how the control strategy adapts to different levels of physical fitness. Although all individuals have been walking all their life and are thus expert walkers, we may still observe different synergies in athletes and non-athletes during perturbated walking. We therefore investigate muscle synergies and movement kinematics while participants walked overground or on a treadmill with either their eyes open or closed. The aim was to investigate how the CNS adapts the gait pattern in these different conditions depending on physical fitness. We hypothesized that athletes and non-athletes share the same muscle synergies but that athletes may recruit additional muscle synergies to reduce coactivation of muscle groups during walking in more challenging conditions.

## 2. Materials and Methods

### 2.1. Experimental Setup

Twenty male participants, ten soccer players (height: 176 ± 5 cm, mass: 72.4 ± 6.8 kg, age: 23 ± 3 years) and ten non-athletes (height: 175 ± 6 cm, mass: 78.0 ± 18.0 kg, age 24 ± 3 years) who were students at the University of Mazandaran, volunteered for the experiment. The athletes had at least seven years continuous training experience and the non-athletes had no exercise training experience during their life. All participants were healthy, right-handed, and did not have any injuries that could affect their gait pattern. Participants were invited to the Centre of Health Assessment and Monitoring of Physical Education at the University of Mazandaran and gave written informed consent prior to beginning the protocol. The procedure was approved by Office of Research Ethics at University of Mazandaran.

Participants were asked to wear comfortable walking shoes and walk overground or on a treadmill (H/P COSMOS treadmill, Germany). To determine the preferred walking speed, we used either the treadmill’s monitor or a stopwatch during 10 m overground walking. To help participants walk on the treadmill with their eyes closed, a rope was applied in the front and back of participants so they could adjust their walking speed when their body touched the rope. After finishing treadmill walking participants rested for 5 min. Then overground walking was performed. A partner was present during eyes-closed overground walking. The partner walked next to the participant while they both held a piece of rope in their hands. Details of walking protocol is provided in figure 1.

**Figure 1.**
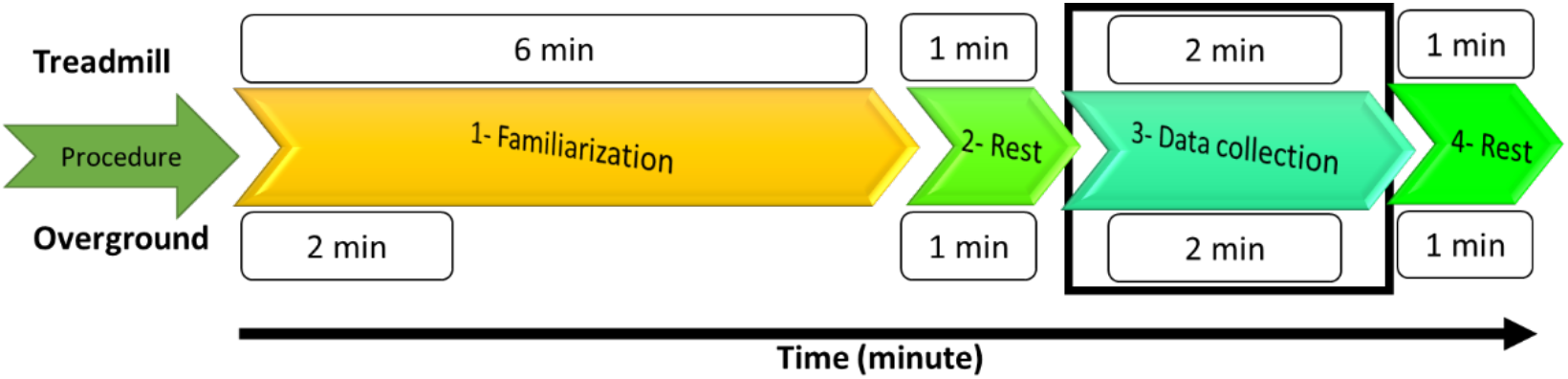
Procedure of treadmill and overground walking. Participants performed treadmill walking before overground walking. The protocol was started with eyes-open condition on each surface. The time duration of treadmill and overground walking during each step are distinguished on the top and down of the procedures, respectively. The black bound box shows when data was collected.

### 2.2. Data Acquisition

During overground walking, lower spinal horizontal rotation was recorded using a 3D inertial measurement unit (IMU) consisting of magnetometers, accelerometers and gyroscopes (Noraxon MyoMotion system, USA). The sensor was placed on lower thoracic (T12). A footswitch was applied to the right leg for the extraction of temporal gait parameters. All kinematic data were sampled at 100 Hz.

Surface EMG was recorded from eight muscles of the right leg (dominant leg, assessed by kicking a ball): vastus medialis (VM), vastus lateralis (VL), biceps femoris (BF), semitendinosus (ST), tibialis anterior (TA), peroneus longus (PL), gastrocnemius medialis (GM), and soleus (S) using a wireless Noraxon myoMuscle system. EMG signals were recorded using a bipolar montage and sampled at 1500 Hz. Kinematic and EEG data were captured synchronously via Noraxon system (analogue input) using the Noraxon MR3.10 analysis software.

### 2.3. Data Preprocessing

Participants walked on a 10-meter walkway during overground walking. They had to turn at the end of walkway and continued their walking for two minutes. We cut data during turning and only analysed data during straight line walking. We used the lower spinal horizontal rotation data to determine the turning points: The zero-crossings indicated the middle of the turning movement and we removed 1.5 s before and after each zero-crossing. The remaining data was used for further analysis. We then segmented the gait cycles using footswitch data for both overground and treadmill walking. Each cycle was defined by consecutive heel strikes and consisted of a stance and swing phase. EMG signals were band-pass filtered at 20-400 Hz (K. M. Steele, Rozumalski, & Schwartz, 2015) and normalised to unit variance. Each gait cycle was then resampled to 1000 data points (0 and 1000 are consecutive heel strikes). In the eyes-closed condition, participants did not always show stable gait dynamics and we removed all gait cycles in which the duration of the gait, stance or swing phase differed more two standard deviation from the mean. EMG signals were then rectified using Hilbert transform to extract the EMG envelopes (Boonstra & Breakspear, 2012; Myers et al., 2003) and low-pass filtered at 15 Hz (Ivanenko, Poppele, & Lacquaniti, 2004).

### 2.4. Extraction of Muscle Synergies

For the extraction of muscle synergies, an equal number of gait cycles was used for each participant and condition. The minimum number of accepted gait cycles was 32 cycles, which were randomly selected from all accepted gait cycles. These gait cycles were concatenated and decomposed using non-negative matrix factorization (NMF). NMF is a linear mode decomposition *X* ↦ *W*^(*m*)^*C*^(*m*)^ that includes the constraints that both extracted activation patterns *C*^(*m*)^ and muscle weighs *W*^(*m*)^ are positive semi-definite, and that *W*^(*m*)^ and *C*^(*m*)^ have rank *m*. We used a multiplicative update algorithm to solve the corresponding minimisation of the Frobenius norm 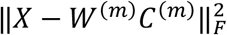 (Lee & Seung, 2001). A set of 1–8 factors (muscle synergies) was iteratively extracted. The reconstruction accuracy was quantified by variance accounted for (VAF): VAF = 1 - SSE/SST, where SSE is the sum of squared errors and SST is the total sum of squares, i.e. the quotient of the Frobenius norm of the error and the Frobenius norm of the rectified EMG, where the error was defined as the difference between the rectified EMG and the product of the muscle weights and temporal activation patterns.

The standard procedure to determine the number of muscle synergies to be extracted is to use the VAF as a fixed threshold (Tang et al., 2015; Torres-Oviedo, Macpherson, & Ting, 2006; Torres-Oviedo & Ting, 2007). Here, we did not use a fixed threshold but varied the threshold based on the individual signal-to-noise ratio. That is, the to-be-found number of synergies will not explain 100% of variance due to noise. Hence, to explain 100% of data variation, spurious muscle synergies will be extracted to explain the noise. However, using a fixed threshold assumes that noise levels are constant across participants and conditions. Instead, we varied the threshold by assessing which of the extracted synergies are robustly expressed across participants and conditions.

To this end, we used k-means clustering to group similar synergies across participants and conditions, as muscle synergies were separately extracted for each participant and experimental condition (Y. Kim, Bulea, & Damiano, 2016; Saltiel, Wyler-Duda, D’Avella, Tresch, & Bizzi, 2001). K-means clustering was performed on the muscle weighs *W*^(*m*)^ of the extracted synergies. To determine the number of clusters to be extracted, we started with one cluster and then sequentially increased the number of clusters until all synergies of the same participant and condition ended up in different clusters. We repeated this procedure for fixed thresholds of 0.7-1 VAF. We expected that the number of clusters would gradually increase with an increasing threshold to group the increasing number of muscle synergies that are extracted. That is, if genuine muscle synergies are extracted, these synergies will be robustly expressed across participants and only a small number of clusters is needed to group them. However, once spurious muscle synergies are extracted, these will vary randomly between participants and conditions and the number of required clusters to group them will rapidly increase. The optimal threshold can then be determined as the point where the number of clusters start to rapidly increase.

Once we determined the number of clusters, i.e., the number of muscle synergies that are robustly expressed across participants and conditions, we further adjusted the threshold on an individual basis. That is, the noise levels may likely vary across participants and conditions, and a variable threshold may therefore be more suitable. We therefore sequentially increased the threshold for individual participants and reran the clustering algorithm. The additional muscle synergies were only kept if the number of clusters needed to group them did not increase.

### 2.5. Module Recruitment Metrics

We computed three metrics to characterise the temporal activation pattern of each muscle synergy (Hayes, Chvatal, French, Ting, & Trumbower, 2014): 1) Module recruitment magnitude (Area) was defined as the area under the curve of recruitment coefficient; 2) Duration of module recruitment (Duty) was defined as the percentage of gait cycle that was above a given threshold; 3) Maximum rate of recruitment coefficient of muscle synergy (Peak). The threshold was set at 15% of maximum recruitment coefficient (Fig. 2). Also, motor module composition was quantified as the sum of the active muscles within a module (sum of weight).

**Figure 2.**
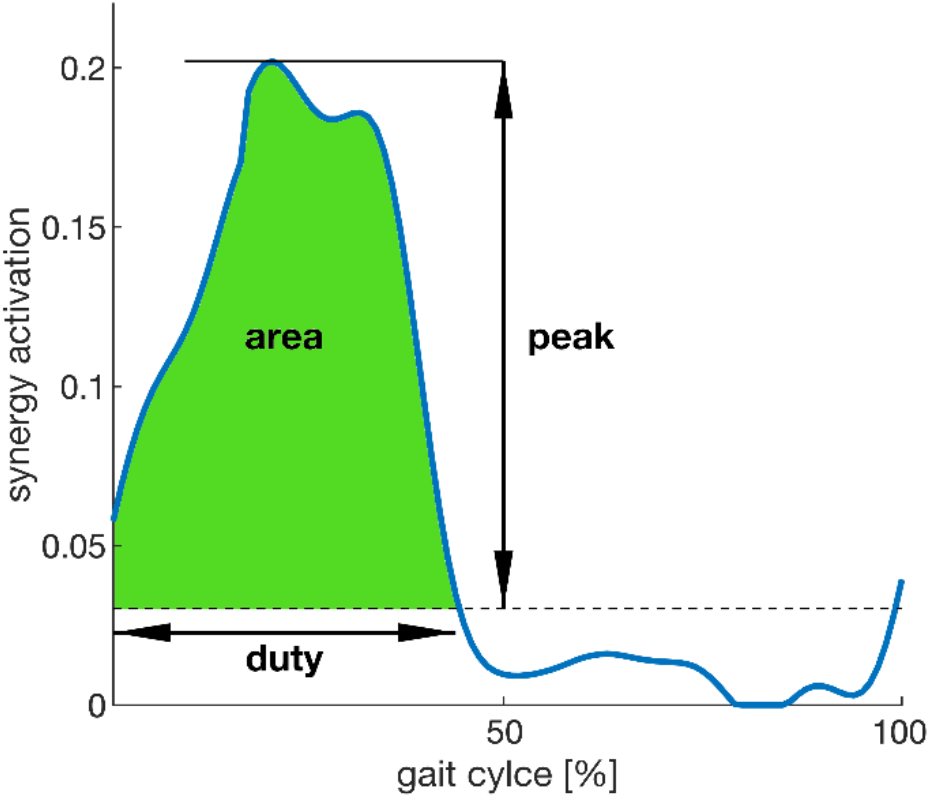
Schematic of synergy activation metrics: Peak is the maximum point of synergy activation, Duty is the duration that synergy activation is greater than the threshold (15% of peak activation), and Area is the area under the mean synergy activation curve and above the threshold.

### 2.6. Statistical Analysis

A linear mixed model was used to compare gait parameters and synergy variables across groups and conditions. In the model we included group (athletes and non-athletes), surface (treadmill and over ground), and vision (eyes-open and eyed-closed walking) as fixed effects factors. As we allowed participants to walk at their preferred walking speed, we also ran the model including walking speed as covariate variable. The Mann-Whitney U test was used to compare the number of synergies between groups, surfaces, and vision conditions respectively. Finally, a binomial generalized mixed effects model was used to compare the expression of specific synergies across groups and conditions. We used Benjamini-Hochberg procedure to adjust p-values for multiple comparisons (Benjamini & Hochberg, 1995). The level of statistical significance was at *α* = 0.05.

## 3. Results

### 3.1. Temporal Gait Parameters

Participants walked slower on treadmill (−48%) compared to overground walking (F_1, 16.2_ = 188.7, p < 0.001) and walked faster with open eyes (26%) compared to closed eyes (F_1, 3.3_ = 77.1, p = 0.04; Fig. 3A). There were no other significant main or interaction effects (p ≥ 0.1). Temporal gait parameters were significantly different between treadmill and overground walking (Fig. 3): Both the cycle and stance duration were longer (a 6% and 13% increase, respectively) during treadmill walking compared to overground walking (F_1, 17.9_ = 11.0, p = 0.04; F_1, 29.6_ = 9.88, p = 0.04). Conversely, swing duration was shorter (−9%) during closed-eyes compared to open-eyes walking (F_1,19.1_ =17.2, p = 0.04). There were no significant main effects for group and no interaction effects for any of the temporal gait parameters (Table S1).

**Figure 3.**
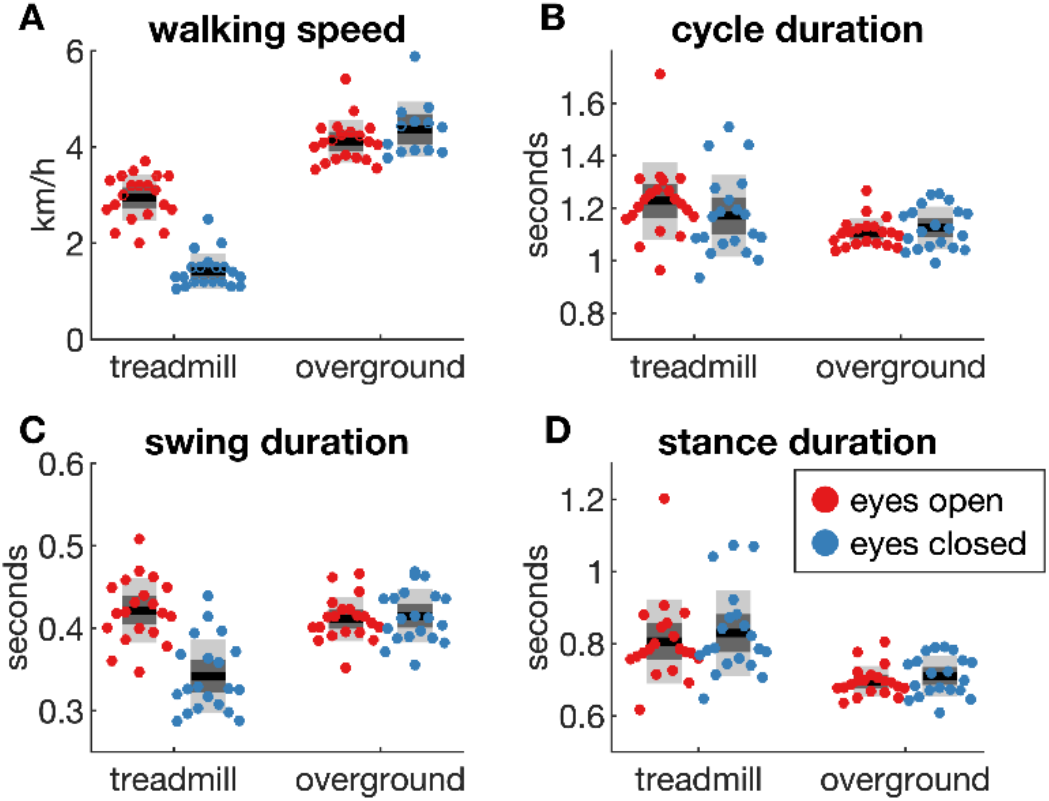
Temporal gait parameters across walking conditions. **A)** Walking speed during eyes-open and eyes-closed walking on treadmill and overground surfaces. Footswitch data was used estimate the average cycle duration (**B**), swing duration (**C**) and the stance duration (**D**). Colored dots show data for individual participants, the black horizontal lines show the group mean, the dark grey box the SEM and the light grey box the SD.

### 3.2. Muscle Synergies

As expected, the explained variances increased with increasing number of synergies: 3 synergies explained about 87% of the variance and 4 synergies 92% (Fig. 4A). We then varied the threshold between 0.7 and 1 and used k-means clustering to group the extracted synergies across participants and conditions. With an increasing threshold, we observed that while the average number of synergies that were extracted in each participant and condition increased gradually, the number of clusters needed to group these synergies showed an abrupt increase from 5 clusters to more than 8 clusters when the threshold was increased above 0.8 (Fig. 4B). These findings show that 5 synergies are robustly expressed across participants and that increasing the threshold above 0.8 results in the extraction of spurious synergies.

**Figure 4.**
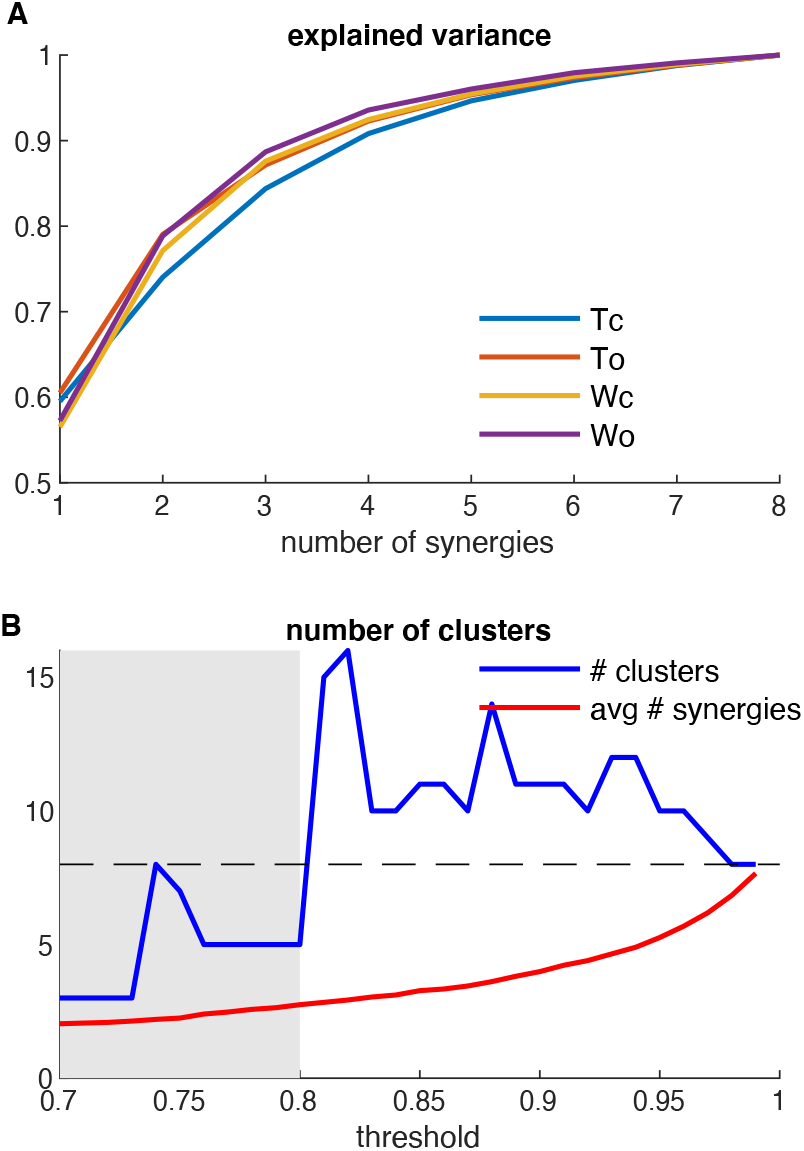
The variance accounted for by each muscle synergy and the number of clusters needed to group them across participants and conditions. **A**) The mean total variance accounted by additional muscle synergies across different experimental conditions. **B**) The average number of synergies extracted for a threshold between 0.7 – 1 (red line) and number of clusters needed to group them (blue line). The horizontal line depicts the number EMGs that were acquired (8 muscles) and signifies the maximum number of synergies that can be extracted.

At the threshold of 0.8, 220 synergies were extracted across participants and conditions (2.8 synergies on average), which explained on average 85.2% of the variance of the EMG envelopes. A fixed threshold may not be optimal in light of potential differences in signal-to-noise ratio across participants and conditions. We therefore sequentially added additional synergies in individual participants and conditions and repeated k-means clustering again after each added synergy, only keeping synergies if it did not increase the number of clusters. This resulted in another 152 synergies that were added. Hence, a total of 372 synergies were extracted across participants and conditions (4.5 synergies on average), explaining 93.7% of the variance, which could be grouped using five clusters. Looking at the muscle weights, most of the synergies remained largely unchanged (e.g., synergies 1, 4, 5), while synergies 2 and 3 revealed a sparser muscle activation pattern (Fig. S1). Using the variable threshold, the five synergies were observed in most of the participants and conditions, allowing for a better statistical comparison across participants and conditions.

The five muscle synergies that were extracted using k-means clustering are shown in figure 5. The first cluster showing BF and ST muscle activation during late stance providing knee flexion during the initial swing phase. The second cluster involves the control of ankle dorsi flexion, with a major involvement of PL muscle and some activation of SO muscle during beginning of stance phase. The third cluster involves activity of the TA muscle and showed two peaks during the stance and swing phase, which can be related to weight bearing during stance and the foot clearance during swing phase, respectively. The fourth cluster showed GM and SO muscle activation during the stance and early swing phase. The fifth cluster involved VM and VL muscles activation during swing phase, providing knee extension important for body weight acceptance before heel contact. Visual comparison suggests that the eyes-open overground walking seems to have the highest peak activation for all clusters. In contrast, activation patterns during eyes-closed treadmill walking appeared more extended and had lower peak activation for all clusters.

**Figure 5.**
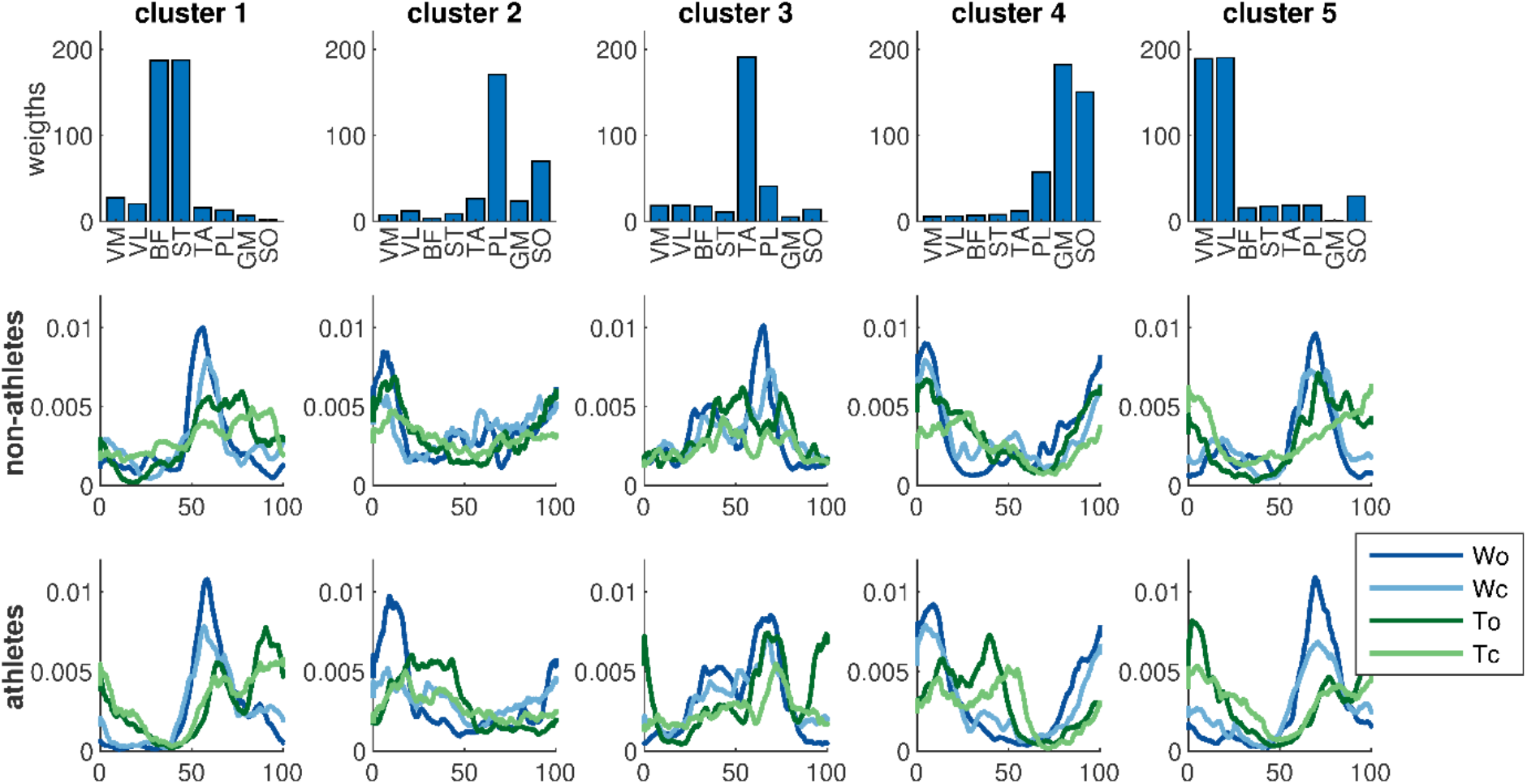
Motor modules extracted from athletes and non-athletes. Activation signals are averaged across participants for both groups during eyes-open and eyes-closed treadmill and overground walking. Conditions are distinguished by colour: WO, overground walking with open eyes; WC, overground walking with closed eyes; TO, treadmill walking with open eyes; TC, treadmill walking with closed eyes.

**Figure 6.**
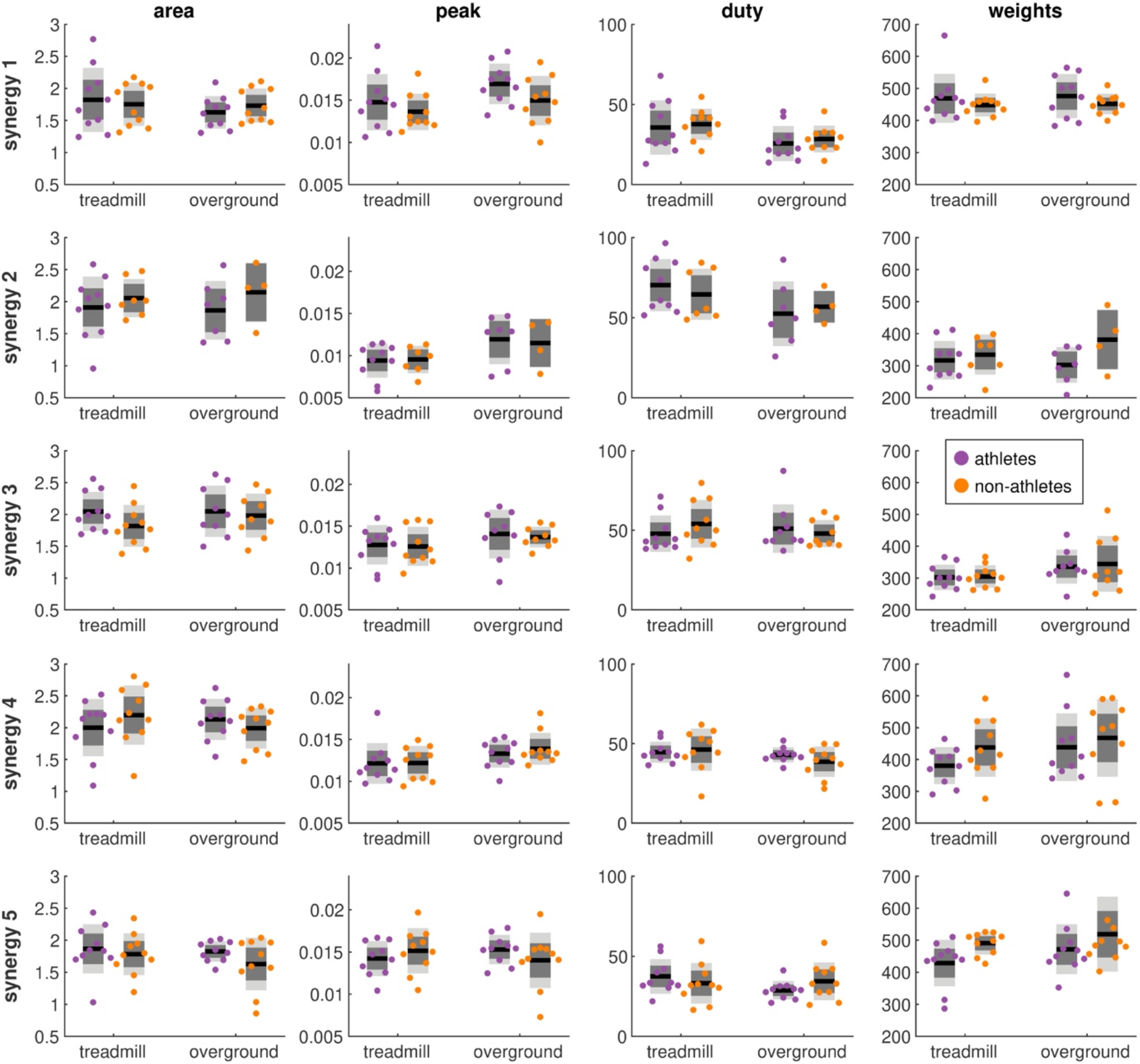
Differences of synergy metrics between groups and walking surface in the eyes-open condition. Rows show the different synergies and column show the different synergy metrics (area, peak, duty and sum of weights). Colored dots show data of individual participants and black horizontal lines show the group mean, the dark grey box the SEM and the light grey box the SD.

### 3.3. Experimental Differences in Muscle Synergies

The number of synergies that were extracted varied between conditions and groups: 192 synergies were extracted during treadmill walking (4.8 ± 0.4 on average) while 180 synergies extracted during overground walking (4.5 ± 0.6), while 186 synergies were extracted during eyes-open walking (4.6 ± 0.6) and 186 during eyes-closed walking (4.6 ± 0.5). Furthermore, 193 synergies were extracted from the athlete group (4.8 ± 0.4) and 179 synergies from the non-athlete group (4.4 ± 0.6). The number of synergies were significantly higher (6%) during treadmill walking than during overground walking (U = 569.5, p = 0.04). Also, athletes showed significantly more synergies (7%) than non-athletes (U = 543.5, p = 0.01). Vision did not have a significant effect on the number of synergies during walking (U = 787, p = 0.93).

We then compared the expression of specific muscle synergies across groups and participants using a binomial generalized mixed effects model. As only synergies 2 and 3 were not recruited in all groups and conditions (Fig. S1), we only tested the recruitment of these two synergies. The model without walking speed as covariate better fitted the data (had smaller information criterion values, see Table S2). We found that synergy 2 was recruited more often in the athlete than in the non-athlete group (87.5% vs. 57.5%, β = −1.9, p = 0.008), irrespective of the addition of walking speed as covariate. No main effects were observed for synergy 3 (p=0.29).

We characterized each muscle synergy using four metrics (area, duty, peak, and sum of weights). We compared these synergy metrics across groups and conditions using a linear mixed model that either included walking speed as covariate or not. In the statistical model without walking speed as covariate a larger number of significant results were found (20 significant effects; Table S3) than with walking speed as covariate (5 significant effects; Table S4). The effects that were significant when including walking speed were also significant when walking speed was not included as covariate, except sum of weight for synergy 4, indicating that these results are statistically robust. We therefore reported the result if they were significant with and without walking speed as covariate. We did not find significant main effects of group for any of these metrics (p > 0.1; Table S4). However, we observed an increased area of synergy 3 and synergy 4 (22% and 18%, respectively) during eyes-open compared to eyes-closed walking. Furthermore, we found increased duty for synergies 5 (26%) during eyes-closed compare to eyes-open walking. We also observed higher peak (14%) during overground walking compare to treadmill walking for synergy 5.

## 4. Discussion

We used muscle synergy analysis to investigate the effect of exercise training on muscle coordination during walking in different gait conditions in athletes and non-athletes. K-means clustering showed that five muscle synergies were robustly expressed across participants and conditions. We found that athletes recruited on average more synergies than non-athletes, that is, they more often recruited synergy 2 consisting of the ankle plantar flexors (PL and SO) that was activated during early stance phase. We also observed some experimental effects across both groups, for example participants recruited additional synergies during treadmill walking compared to overground walking. In addition, the area of the activity pattern of synergy 3 and synergy 4 was significantly reduced during the eyes-closed compared to eyes-open condition, whereas duty of synergy 5 was significantly increased during the eyes-closed compared to eyes-open condition. Further, peak of activation was greater during overground walking compared to treadmill walking for synergy 5. These results suggest that exercise training results in subtle differences in muscle coordination during gait that can be detected using synergy analysis.

Synergy analysis showed that athletes used a greater number of synergies than non-athletes (4.85 vs. 4.48). These findings are in line with previous literature suggesting that athletes generally recruit more synergies (Sawers et al., 2015) while patients recruit less synergies (Tang et al., 2015). Recruiting additional synergies provides greater flexibility of muscle coordination and may reduce muscle coactivation, although caution should be exerted when interpreting synergy number without comprehensive assessment of other synergy metrics (Hayes et al., 2014; Sawers et al., 2015). In addition, when using a fixed threshold to extract synergies, it is possible that a change in synergy number is caused by relative changes in the VAF of individual synergies. Here we used a variable threshold and k-means clustering to determine whether each synergy was expressed in individual participants and conditions. This approach also allowed to determine the specific synergy that was additionally recruited. We found that athletes more often recruited synergy 2 involving the ankle plantar flexors (PL and SO muscles) than non-athletes. Computation modelling reveals that the increased activation of the plantar flexors results in a shift from ‘heel-walking’ to ‘toe-walking’ gait (Ong, Geijtenbeek, Hicks, & Delp, 2019). In parallel with the previous study, it is likely that soccer players with more than 7 years of exercise training have greater plantar flexor strength (Fousekis, Tsepis, & Vagenas, 2010) and are therefore more likely to recruit this synergy during walking.

In addition to synergy number, we also assessed four synergy metrics. Contrary to our expectations, we did no find significant group differences for any of the synergy metrics. For example, athletes and non-athletes showed similar sum of weights for all synergies extracted during treadmill and overground walking (Table S4). This is in contrast to a previous study that showed that athletes had reduced muscle weights during overground and beam walking (Sawers et al., 2015). They compared professional trained ballet dancers with novices (no dance or gymnastics training) and reported that professional dancers had a higher rate of spatial orientation because of their superior orientation and position in the space, which corresponded to training experience and proper coordination (Crotts, Thompson, Nahom, Ryan, & Newton, 1996). The effect of exercise training on muscle coordination may hence depend on the field of sport.

Although we did not observe any difference between group, there were several experimental changes across surface or vision condition. Our results showed more extended duty of synergy 5 (involving the VM and VL muscles) during overground compared to treadmill walking. Knee extensors play an important role in early stance of gait cycle by supporting body weight and controlling knee extension (Arnold, Anderson, Pandy, & Delp, 2005; Siegel, Kepple, & Stanhope, 2006). The strength of these muscles can help individuals with a crouch gait to walk in a more upright posture (K. Steele, Damiano, Eek, Unger, & Delp, 2012). There are controversial outcomes of knee joint function during walking on treadmill and overground surfaces. Extended treadmill walking (more than 6 min) may create a shift from visual to leg-proprioceptive gait control (Prokop, Schubert, & Berger, 1997). However, it has been reported that kinematics of knee during treadmill walking was similar to overground walking (Wass, Taylor, & Matsas, 2005). Conversely, VL muscle had shorter duration, later onset, and earlier offset when participants walked on a treadmill. Treadmill walking caused immediate changes in VL muscle too (Harris-Love, Macko, Whitall, & Forrester, 2004). As stance phase can increase during treadmill walking (Chockalingam, Chatterley, Healy, Greenhalgh, & Branthwaite, 2012) and participants may generate a more extended duty of VM and VL muscles to provide more optimal shock absorption during loading phase when walked on treadmill (Siegel et al., 2006).

In addition, our results showed that visual information can have an effect on synergy metrics. We observed reduced activation (smaller area) of synergy 3 (TA muscle) and synergy 4 (GM and SO muscles) during eyes-closed walking. Weakness of TA muscle can dramatically decrease foot stability (Gefen, 2001). TA normally active during loading response and pre-swing phases to prepare the foot for initial contact and decelerate the rate of foot fall during swing phase, respectively (Byrne, O’keeffe, Donnelly, & Lyons, 2007). Also, GM and SO muscles are crucial to add more energy to the trunk during stance phase (Riley, Della Croce, & Kerrigan, 2001; Siegel, Kepple, & Stanhope, 2004) and provide ankle stability during gait (Sutherland, Cooper, & Daniel, 1980). Weakness of plantar flexor muscles can dramatically decrease foot stability and lead to fall risk (Cattagni, Scaglioni, Laroche, Grémeaux, & Martin, 2016). Plantar flexor muscles are strongly required to support weight of body and provide stability at the ankle joint during gait (Cattagni et al., 2018). Visual information has a fundamental role in the CNS circuits to perform skilled movement, precise visuomotor coordination is need for accurate gait (Georgopoulos & Grillner, 1989). Visual information is registered and stored in short term-memory as a "body schema" which encompass the body position in space and body parts associated with movement (Massion, 1992). It seems when participants closed their eyes this ability of visuomotor system dramatically reduced and participants may be less able to predict the moment of heel strike, with lower activity of TA, GM and SO muscles (ankle stabilizers), during eyes-closed walking to create a safer gait.

In addition to changes in muscle coordination, we also found changes in temporal gait parameters. During treadmill walking the cycle and stance duration increased, which confirms a previous study that examined the effect of treadmill and overground walking on gait kinematics (Malatesta, Canepa, & Fernandez, 2017). Participants may increase cycle and stance duration during treadmill walking because they were less familiar with treadmill walking (Nagano, Begg, Sparrow, & Taylor, 2013; Watt et al., 2010). Furthermore, a trend of shorter swing phase during treadmill compared to overground walking was reported, although they found no significant difference in kinematics measurements (Murray, Spurr, Sepic, Gardner, & Mollinger, 1985). Interestingly, our results showed a decreased swing duration during eyes-closed compare to eyes-open walking. It can help to increase gait stability in absence of visual information. The absence of difference in cycle or stance duration during different visual conditions, suggest that decreased in swing duration during eyes-closed walking results from the decreased in actual swing duration (Layne et al., 2018).

We allowed participants to walk at their preferred walking speed in both conditions, which resulted in slower walking speed during treadmill compared to overground walking and during eyes-closed compared eyes-open walking. To test for potential confounding effects of walking speed on muscle synergies, we included walking speed as covariate in our statistical analyses. This indeed reveals that some experimental effects are mediated through a change in walking speed (see Table S3 and S4). However, the group effect on the recruitment of synergy 2 was unaffected by including walking speed as covariate. Walking speed has been shown to affect muscle coordination during gait (Kibushi, Hagio, Moritani, & Kouzaki, 2018) and it may be better to control it in future studies, although this may not be easily achieved in an eyes-closed condition. Finally, the current study had a fairly limited sample size (n=10 in each group) and the reported effects of exercise training on muscle coordination in gait should therefore be replicated in larger samples.

## 4. Conclusion

This paper aimed to investigate the effect of long-term exercise training by comparing athlete and non-athlete subjects during treadmill and overground walking. Our results indicated that synergy analysis is able to detect different neuromuscular strategies in athletes and non-athletes. Athletes recruit more muscle synergies than non-athletes, in particular they more often recruit the plantar flexor synergy. In conclusion, the differences in athletes were observed in the coordination of the ankle joint stabilizers that may help to create a more flexible and stable gait.

## Acknowledgments

We would like to thank the volunteer participants for their participation in the experiments and also the Otech_motionlab group for their cooperation. TB has received funding from the European Union’s Horizon 2020 research and innovation programme under the Marie Sklodowska-Curie grant agreement No 895914.

## Conflict of interest

The authors declare that they have no conflict of interest.

## Notes

### Competing Interest Statement

The authors have declared no competing interest.

