## Supplemental Material for "The effects of long-term exercise training on the neural control of walking"

**Table S1. Statistics analysis of temporal gait parameters and walking speed**

| | Group | | Surface | | Vision | | Group*vision | | Group*surface | |
|---|---|---|---|---|---|---|---|---|---|---|
| | F | $P_{adj}$* | F | $P_{adj}$* | F | $P_{adj}$* | F | $P_{adj}$* | F | $P_{adj}$* |
| Swing duration | 0.37 | 0.88 | 3.23 | 0.36 | **17.27** | **0.04** | 0.98 | 0.71 | 0.04 | 0.99 |
| Stance duration | 0.92 | 0.72 | **9.88** | **0.04** | 1.73 | 0.63 | 5.54 | 0.22 | 0.14 | 0.95 |
| Cycle duration | 1.12 | 0.71 | **10.98** | **0.04** | 2.07 | 0.57 | 0.53 | 0.84 | 0.50 | 0.84 |
| Walking speed | 7.31 | 0.13 | **188.77** | **<0.001** | **77.10** | **0.04** | 0.01 | 0.99 | 0.64 | 0.82 |

Significant effects (P<0.05) are displayed in bold. * P-values are adjusted using the Benjamini-Hochberg procedure.

**Table S2. Statistical results of a binomial generalized mixed effects model of the recruitment of muscle synergies**

| | **Synergy 2** | | | | **Synergy 3** | | | |
|---|---|---|---|---|---|---|---|---|
| **Model fit** | No covariate | | With covariate* | | No covariate | | With covariate* | |
| AIC | 26.4 | | 68.4 | | 22.9 | | 46.0 | |
| BIC | 35.4 | | 79.0 | | 31.9 | | 56.5 | |
| **Fixed effects** | β | P | β | P | β | P | β | P |
| Group | **-1.9** | **0.008** | **-2.2** | **0.002** | -0.76 | 0.43 | -0.59 | 0.52 |
| Surface | **-1.9** | **0.008** | -1.6 | 0.15 | -0.76 | 0.43 | -2.1 | 0.36 |
| Vision | 0.33 | 0.40 | 0.11 | 0.85 | -0.76 | 0.43 | -0.99 | 0.29 |

Significant effects (P<0.05) are displayed in bold. * Walking speed is added as covariate. Only synergies 2 and 3 were analysed as these were not expressed in all participants and conditions.

**Table S3. Statistical results of the linear mixed model of the muscle synergy metrics without walking speed as covariate variable**

| | | Group effect | | Surface effect | | Vision effect | | Group*vision | | Group*surface | |
|---|---|---|---|---|---|---|---|---|---|---|---|
| | | F | $P_{adj}$* | F | $P_{adj}$* | F | $P_{adj}$* | F | $P_{adj}$* | F | $P_{adj}$* |
| Synergy 1 | Area | 0.00 | 0.98 | 5.27 | 0.118 | 0.95 | 0.58 | 0.49 | 0.71 | 1.19 | 0.58 |
| | Peak | 1.83 | 0.44 | **40.28** | **<0.001** | **11.29** | **0.02** | 0.48 | 0.71 | 0.21 | 0.83 |
| | Duty | 0.18 | 0.84 | **63.87** | **<0.001** | 0.34 | 0.77 | 0.08 | 0.92 | 0.01 | 0.97 |
| | Sum | 0.89 | 0.58 | 7.85 | 0.06 | 0.00 | 0.98 | 0.28 | 0.79 | 1.28 | 0.57 |
| Synergy 2 | Area | 3.33 | 0.23 | 8.17 | 0.09 | 0.06 | 0.93 | 0.46 | 0.77 | 4.93 | 0.17 |
| | Peak | 0.00 | 0.98 | 7.37 | 0.09 | **10.80** | **0.03** | 0.00 | 0.99 | 0.05 | 0.93 |
| | Duty | 0.21 | 0.83 | 8.74 | 0.06 | **9.43** | **0.04** | 0.93 | 0.58 | 1.14 | 0.58 |
| | Sum | 3.62 | 0.22 | 0.98 | 0.58 | 1.11 | 0.58 | 0.29 | 0.79 | 0.94 | 0.58 |
| Synergy 3 | Area | 0.62 | 0.66 | **10.42** | **0.03** | **21.88** | **0.002** | 0.03 | 0.94 | 5.00 | 0.13 |
| | Peak | 0.03 | 0.95 | **22.31** | **<0.001** | **8.57** | **0.047** | 0.84 | 0.59 | 0.75 | 0.61 |
| | Duty | 0.92 | 0.58 | 2.92 | 0.27 | 1.70 | 0.458 | 2.13 | 0.40 | 1.33 | 0.57 |
| | Sum | 0.92 | 0.58 | **14.06** | **0.02** | 1.95 | 0.423 | 0.06 | 0.93 | 3.14 | 0.26 |
| Synergy 4 | Area | 0.08 | 0.92 | 5.26 | 0.12 | **24.82** | **0.001** | 0.24 | 0.81 | 0.34 | 0.77 |
| | Peak | 0.71 | 0.62 | **31.06** | **<0.001** | 0.32 | 0.77 | 0.01 | 0.98 | 7.00 | 0.07 |
| | Duty | 0.45 | 0.73 | **22.80** | **0.001** | 0.05 | 0.93 | 2.49 | 0.33 | **10.55** | **0.03** |
| | Sum | 3.44 | 0.22 | 6.31 | 0.09 | 0.35 | 0.77 | 0.01 | 0.97 | 0.03 | 0.94 |
| Synergy 5 | Area | 0.49 | 0.71 | 7.02 | 0.07 | 4.34 | 0.16 | 1.91 | 0.42 | 0.10 | 0.92 |
| | Peak | 0.18 | 0.84 | **41.55** | **<0.001** | **18.30** | **0.004** | 1.68 | 0.46 | **9.76** | **0.03** |
| | Duty | 0.09 | 0.92 | **53.23** | **<0.001** | **10.13** | **0.03** | 0.94 | 0.58 | **12.83** | **0.02** |
| | Sum | 5.21 | 0.12 | 5.71 | 0.10 | 1.08 | 0.58 | 1.18 | 0.58 | 0.01 | 0.97 |

Significant effects (P<0.05) are displayed in bold. * P-values are adjusted using the Benjamini-Hochberg procedure

**Table S4. Statistical results of the linear mixed model of the muscle synergy metrics with walking speed as covariate variable**

| | | Group effect | | Surface effect | | Vision effect | | Group*vision | | Group*surface | |
|---|---|---|---|---|---|---|---|---|---|---|---|
| | | F | $P_{adj}$* | F | $P_{adj}$* | F | $P_{adj}$* | F | $P_{adj}$* | F | $P_{adj}$* |
| Synergy 1 | Area | 0.01 | 0.99 | 0.25 | 0.90 | 0.09 | 0.97 | 0.01 | 0.99 | 1.72 | 0.63 |
| | Peak | 1.18 | 0.71 | 0.31 | 0.88 | 0.02 | 0.99 | 0.00 | 0.99 | 2.83 | 0.42 |
| | Duty | 0.00 | 0.99 | 0.17 | 0.94 | 0.04 | 0.99 | 0.03 | 0.99 | 0.34 | 0.88 |
| | Sum | 0.61 | 0.82 | 0.01 | 0.99 | 0.23 | 0.90 | 0.03 | 0.99 | 3.93 | 0.33 |
| Synergy 2 | Area | - | - | - | - | - | - | - | - | - | - |
| | Peak | - | - | - | - | - | - | - | - | - | - |
| | Duty | 0.70 | 0.81 | 0.04 | 0.99 | 1.17 | 0.71 | 0.25 | 0.90 | 0.19 | 0.94 |
| | Sum | - | - | - | - | - | - | - | - | - | - |
| Synergy 3 | Area | 0.11 | 0.97 | 0.58 | 0.82 | **18.05** | **0.04** | 1.35 | 0.71 | 5.55 | 0.22 |
| | Peak | 0.33 | 0.88 | 5.27 | 0.22 | 1.67 | 0.64 | 0.01 | 0.99 | 0.29 | 0.90 |
| | Duty | 0.36 | 0.88 | 2.34 | 0.46 | 2.83 | 0.43 | 3.68 | 0.36 | 1.22 | 0.71 |
| | Sum | 1.04 | 0.71 | 0.60 | 0.82 | 0.35 | 0.88 | 0.00 | 0.99 | 3.11 | 0.43 |
| Synergy 4 | Area | 0.02 | 0.99 | 0.02 | 0.99 | **19.17** | **0.04** | 3.68 | 0.36 | 0.07 | 0.99 |
| | Peak | 0.56 | 0.83 | 0.16 | 0.94 | 0.13 | 0.96 | 3.81 | 0.36 | 1.03 | 0.71 |
| | Duty | 0.10 | 0.97 | 1.23 | 0.71 | 0.00 | 0.99 | 5.16 | 0.22 | 5.58 | 0.20 |
| | Sum | 1.68 | 0.63 | **9.49** | **0.04** | 3.12 | 0.36 | 0.16 | 0.94 | 0.03 | 0.99 |
| Synergy 5 | Area | 0.92 | 0.72 | 0.01 | 0.99 | 1.22 | 0.71 | 1.45 | 0.71 | 0.04 | 0.99 |
| | Peak | 1.01 | 0.71 | **11.32** | **0.04** | 7.60 | 0.15 | 3.02 | 0.42 | 6.25 | 0.19 |
| | Duty | 0.71 | 0.81 | 0.36 | 0.88 | **12.72** | **0.04** | 2.74 | 0.45 | 8.20 | 0.12 |
| | Sum | 4.81 | 0.22 | 0.89 | 0.72 | 0.27 | 0.90 | 0.44 | 0.88 | 0.00 | 0.99 |

Significant effects ($P<0.05$) are displayed in bold. * P-values are adjusted using the Benjamini-Hochberg procedure. The model did not converge for synergy 2 most likely due to missing values (synergy 2 was not recruited in all participants and conditions).

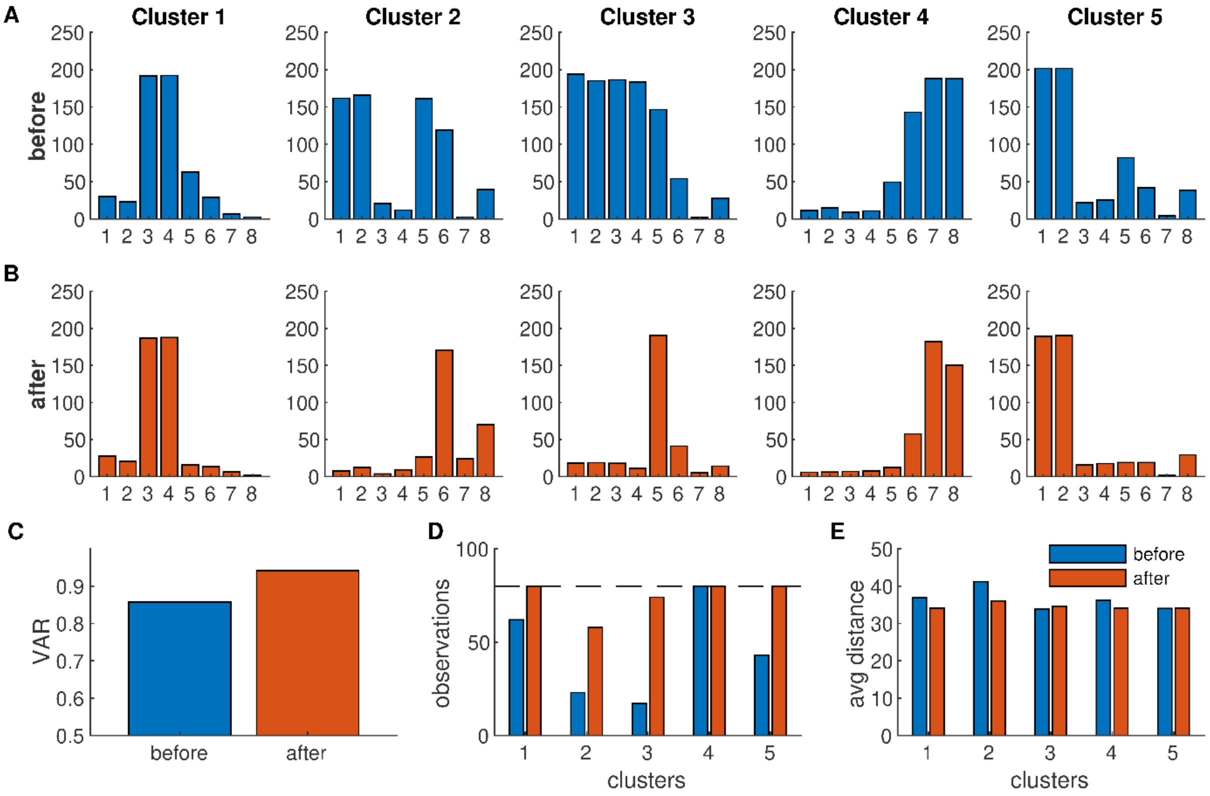


**Figure S1**: **Muscle synergies extracted with fixed and variable threshold**. **A)** The muscle weights of the 5 clusters extracted with a fixed threshold (80% VAF). Synergies are extracted separately for each participant and condition and then grouped using k-means clustering such that all synergies from the same participant and condition end up in different clusters. The muscle weights reflect the centroid of each cluster. **B)** The muscle synergies extracted with a variable threshold. Additional synergies are sequentially added, and k-means clustering is performed again after each added synergy. The added synergy is kept if the number of clusters remains unchanged or removed otherwise. **C)** The explained variance is increased from 85.2% to 93.7% when changing from a fixed to a variable threshold. **D)** The number of observations in each cluster. Using a variable threshold, the 5 synergies are observed in most of the participants and conditions (only synergies 2 and 3 are not observed in all participants and conditions). The dashed line reflects the maximum number of observations (20 participants $\times$ 4 conditions = 80 observations). **E)** The average distance between each observation and the centroid of the corresponding cluster.
